# The CENP-A chaperone complex spatially organizes centromeres

**DOI:** 10.1101/2025.11.07.687291

**Authors:** Hindol Gupta, Julian Haase, Yicong Wu, Jiji Chen, George Karadimov, Hari Shroff, Alexander E. Kelly

## Abstract

Centromeres, defined by CENP-A-containing nucleosomes, direct the assembly of kinetochores for spindle attachment. In mitosis, CENP-A and the constitutive centromere-associated network (CCAN) of the inner kinetochore are arranged into bipartite subdomains within clearings of chromatin. However, it remains unclear whether any of these features exist before mitosis. We show that in interphase, CENP-A and the CCAN assemble ∼200-300 nm shell-like structures that enclose a chromatin-poor central cavity. Strikingly, this cavity is occupied by the interphase-specific CENP-A chaperone complex, which promotes CENP-A assembly once per cell cycle. However, chaperone presence, but not CENP-A incorporation, is required to generate both the shell architecture and the chromatin clearing. The CCAN scaffold CENP-C, which links CENP-A nucleosomes to the chaperone complex, exhibits radial organization spanning the entire structure and is essential for its formation. These data uncover a previously unrecognized structural role for the CENP-A chaperone machinery in establishing interphase centromere architecture and suggest a mechanism by which this machinery configures centromeres for faithful kinetochore assembly and genome stability.

## Introduction

Centromeres are specialized chromosomal loci that direct assembly of the kinetochore, which mediates spindle attachment to ensure accurate chromosome segregation (*1*). Centromeres are specified by nucleosomes containing the histone H3 variant CENP-A, which recruits the constitutive centromere-associated network (CCAN) proteins to build the inner kinetochore (*2*). In mitosis, the CCAN recruits the outer kinetochore, which directly interacts with spindle microtubules. Centromeres are epigenetically maintained by the CENP-A chaperone complex consisting of HJURP and its cofactors, Mis18α, Mis18β, and Mis18BP1 (*3*). During interphase in most eukaryotes, the CENP-A chaperone machinery assembles new CENP-A nucleosomes to restore CENP-A levels after dilution due to DNA replication in the previous cell cycle. The CCAN component CENP-C directs the CENP-A chaperone complex to sites containing pre-assembled CENP-A nucleosomes (*4–8*), a process regulated by kinases Polo-like kinase 1 (PLK1) and CDK1 (*5*, *9–15*). Thus, the CCAN is important for CENP-A assembly in interphase and for the assembly of the complete kinetochore in mitosis.

Within the core centromere, CENP-A nucleosomes are interspersed with canonical histone H3 nucleosomes (*16*), with only ∼100 CENP-A nucleosomes on mitotic chromosomes in human cells (*17*). Despite this low density, CENP-A nucleosomes are closely organized in three dimensions on the surface of mitotic chromosomes, likely to promote the proper capture and attachment of spindle microtubules (*18*). How this three-dimensional arrangement is manifested is a major question in biology. Recent work has revealed the mitotic centromere organization in greater detail. Individual centromeres and kinetochores are partitioned into two subdomains by condensin and cohesin (*19*). Further, cryo-ET reconstructions of intact human mitotic centromeres revealed that each subdomain resides within a chromatin-clearing, distinct from bulk chromatin (*20*). These clearings consist of ∼250 nucleosomes, of which about 50, presumably containing CENP-A, are directly engaged with the CCAN. CENP-C is required to maintain these clearings, which are thought to concentrate CCAN complexes to promote kinetochore formation and microtubule attachment. However, it remains unclear how these clearings arise and whether they form prior to or during mitosis. Super-resolution imaging in human cells has visualized a G1-specific rosette-like arrangement of CENP-A immediately following HJURP-mediated deposition (*21*), suggesting that a specific centromere arrangement exists prior to mitosis.

In this study, we investigated the *de novo* assembly of centromere and kinetochore architecture in *Xenopus* embryo extracts using advanced super-resolution imaging and deep learning resolution enhancement. We find that CENP-A and CCAN subunits assemble ∼200–300 nm shell-like structures in interphase that enclose a chromatin-poor central cavity, which is occupied by the CENP-A chaperone complex. Chaperone presence, but not CENP-A incorporation, is required to generate both the shells and the chromatin clearing, and the CCAN scaffold CENP-C is radially oriented to connect CENP-A chromatin to the chaperone core and is essential for structure formation. These results reveal a previously unrecognized structural role for the CENP-A chaperone machinery in establishing interphase centromere architecture and suggest a mechanism by which this machinery preconfigures centromeres and kinetochores to prime their organization in mitosis.

## Results

### The centromere and inner kinetochore form hollow shell-like structures during interphase

In mitosis, the CCAN and its associated CENP-A nucleosomes have a defined three-dimensional arrangement within clearings of chromatin that is thought to promote proper kinetochore-microtubule attachment (*19*, *20*). However, it remains unclear how the CCAN and CENP-A are arranged during interphase, and whether there is a unifying principle governing centromere architecture throughout the cell cycle. To explore this, we imaged the CCAN protein CENP-T by 3D-SIM super-resolution microscopy at different time points in *Xenopus* egg extracts, which are amenable to strict cell-cycle control (Fig. 1A and Figs. S1A and S1B). CENP-T formed puncta in early interphase, forty minutes after entering interphase, but formed ring-like structures later in interphase, when replication is complete. CENP-T rings were smaller but still observable ten minutes after entry into M phase. However, they were no longer resolvable at metaphase, when sister-chromatids are fully separated (Fig. 1A and Figs. S1A and S1B). CENP-C also transitioned from a punctate to a ring-like structure during interphase (Fig. S1C). These ring-like structures were consistently observed at nearly all kinetochores (Fig. S1D) and were detectable using distinct super-resolution technologies (Fig. S1E). Interestingly, the formation of CENP-C and CENP-T rings in interphase was not dependent on replication (Fig. S1F-I), indicating that the fundamental architecture of individual inner kinetochores in interphase is ring-like. Thus, our results demonstrate that the spatial organization of the CCAN is modulated throughout the cell cycle, independently of DNA replication.

**Fig. 1.**
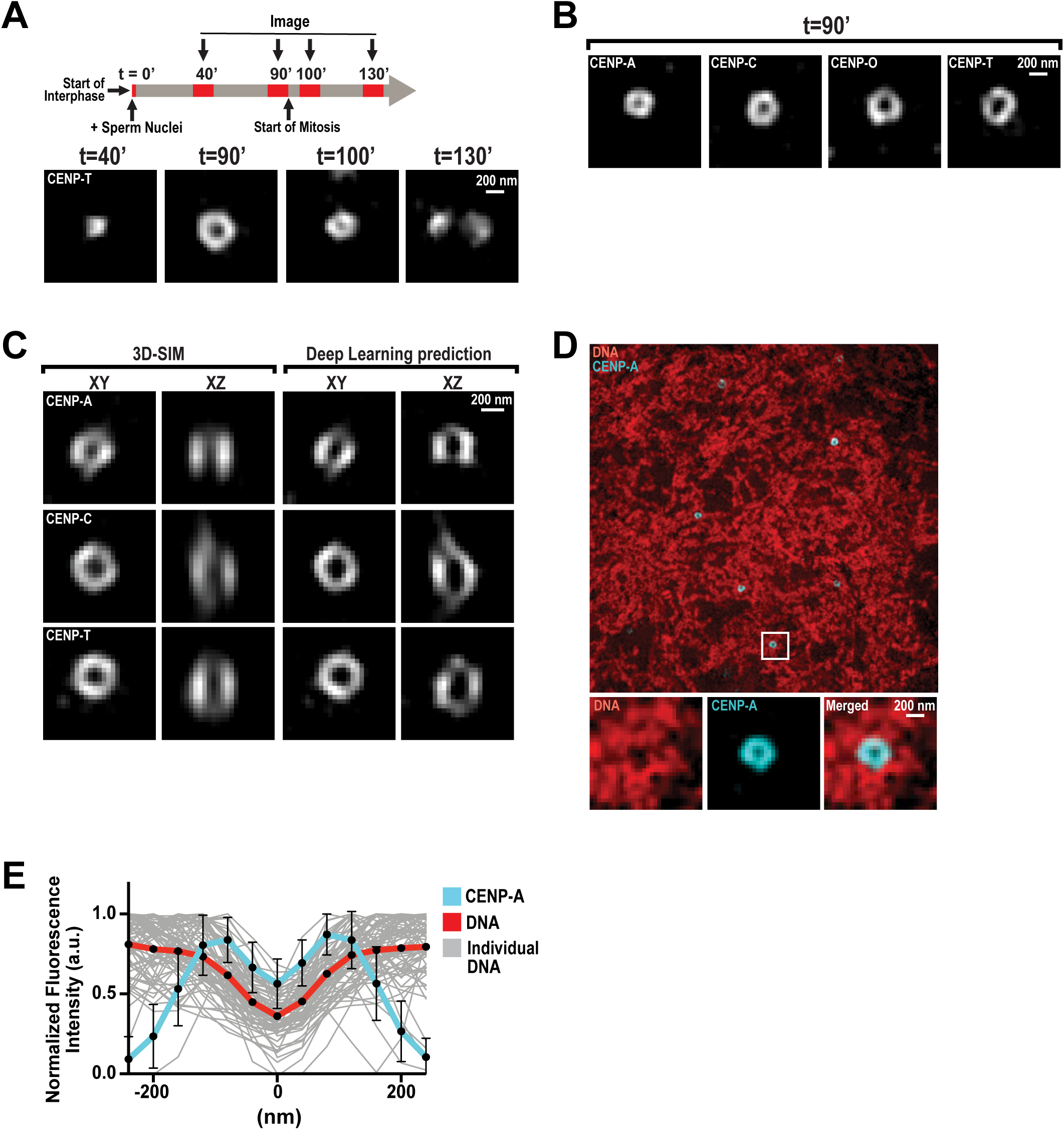
The centromere and inner kinetochore form shell-like structures around chromatin clearings in interphase. (**A**) Upper: Experimental timeline for imaging kinetochores on interphase nuclei and mitotic chromosomes in *Xenopus* egg extracts. Samples were taken at the indicated times (in minutes) after the addition of sperm and calcium, and processed for immunofluorescence staining. Lower: Representative 3D-SIM immunofluorescence images of CENP-T at indicated timepoints. (**B**) Representative 3D-SIM images of CENP-A, CENP-C, CENP-O, and CENP-T on interphase nuclei, at *t*=90’. Scale bar is 200 nm. (**C**) Lateral and axial views of representative 3D-SIM images of CENP-A, CENP-C, and CENP-T on interphase nuclei before and after axial resolution enhancement by deep learning analysis. Scale bar is 200 nm. (**D**) Representative 3D-SIM image of CENP-A (cyan) and DNA (DAPI; red) on interphase nuclei, at *t*=90’. Higher magnification views of the white box region below, showing individual and merged fluorescence channels. Scale bar is 200 nm. (**E**) Linescan quantification of CENP-A and DNA fluorescence at centromeres from (D). Bold lines indicate average intensity, and thin gray lines indicate individual traces of DNA fluorescence. Each trace was aligned by the central minimum and normalized to the maximum intensity. n=75 centromeres. Error bars represent standard deviation (SD).

We found that CENP-A and the CCAN protein CENP-O also formed stereotypical ring-like structures in interphase (Fig. 1B). Measurement of the average ring diameter of CENP-A and CCAN structures revealed a hierarchy of nested rings wherein CENP-A and CENP-C rings had the smallest average diameter (191 ± 30 and 227 ± 33 nm, respectively), and CENP-O and CENP-T the largest (267 ± 43 and 278 ± 42 nm, respectively) (Fig. S2A). These data suggest that the CCAN is assembled around CENP-A rings, with the spatial arrangement of CCAN subunits relative to CENP-A in agreement with cryo-EM studies on single CCAN-CENP-A-nucleosome complexes (*22*, *23*). Next, we sought to determine the three-dimensional arrangement of CENP-A and CCAN in interphase nuclei. Although our 3D-SIM imaging of kinetochores achieved lateral resolutions of ∼125 nm, the axial resolution was about three times poorer (∼340-380 nm). Therefore, we used a deep learning approach that yields isotropic 120-nm resolution on our 3D-SIM images of CENP-A and the CCAN (*24*). This approach predicts that CENP-A, CENP-C, and CENP-T form ring-like structures in both lateral and axial slices at the centroid of each structure (Fig. 1C). However, the arrangement of the signals in the axial dimension is less regular and slightly oblong when compared to the lateral slices. These data indicate that CENP-A and the CCAN form hollow, semi-enclosed particles in interphase.

Although CENP-A-containing chromatin is not present within these structures, canonical histone H3-containing chromatin may be. To test this, we co-imaged DNA and CENP-A using 3D-SIM (Fig. 1D). Remarkably, the levels of DNA were significantly reduced within the central cavity of the CENP-A structures, as indicated by quantitative line scans (Fig. 1E). In addition, linescan analysis suggested that chromatin density begins to decrease at the outer diameter of the CENP-A ring-like structure. To further examine the relative location of DNA and CENP-A, we processed the 3D-SIM images using the deep-learning protocol (Fig. 2B). In resolution-enhanced images, there is an apparent decrease in DNA staining within the CENP-A shell and the central cavity. These data suggest a large clearing of chromatin, with CENP-A shell-like structures defining its borders in three dimensions.

**Fig. 2.**
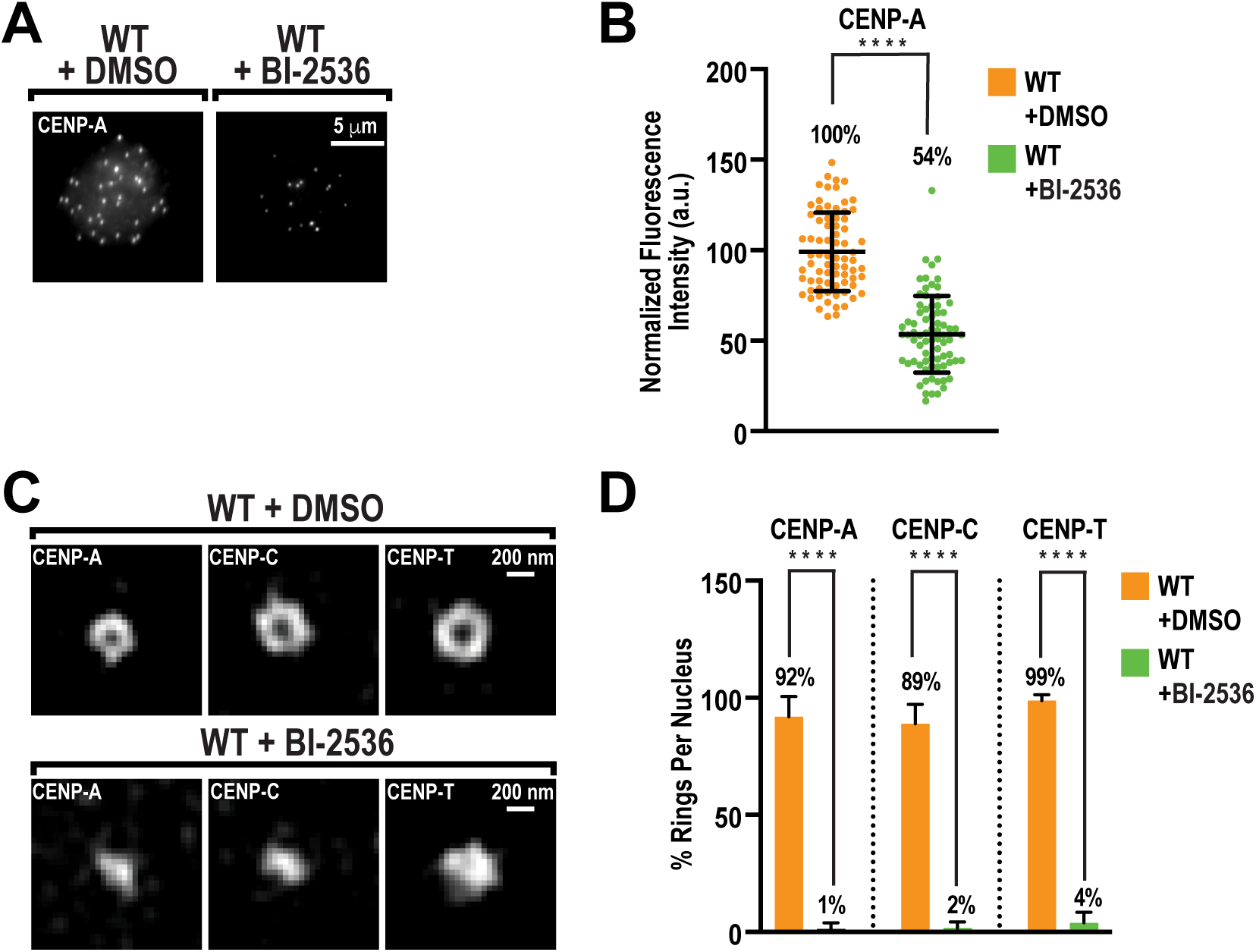
PLK1 inhibition perturbs centromere and kinetochore architecture. (**A**) Representative immunofluorescence images of interphase nuclei stained for CENP-A, at *t*=90’, in extract treated with DMSO or BI-2536. Scale bar is 5 µm. (**B**) Quantification of fluorescence intensity from (A), normalized to DMSO. n = 75 centromeres per condition. A.U., arbitrary units. Error bars represent SD, ****, *P* < 0.0001. (**C**) Representative 3D-SIM images of CENP-A, CENP-C, and CENP-T on interphase nuclei, at *t*=90’, in extract treated with DMSO or BI-2536. Scale bar is 200 nm. (**D**) Percentage of observable ring-like structures per nucleus for conditions in (C). n=10 nuclei per condition, and nβ145 centromeres total. Error bars represent SD, ****, *P* < 0.0001.

### PLK1 activity is required for centromere architecture

During interphase in vertebrate cells, new CENP-A molecules are assembled at centromeres by the CENP-A chaperone complex to preserve centromere identity (*3*). The chaperone HJURP, along with the Mis18 complex (Mis18α, Mis18β, and Mis18BP1), constitutes the chaperone complex and promotes the assembly of CENP-A into centromeric chromatin to replenish CENP-A levels after dilution due to DNA replication from the prior cell cycle. Previous work has demonstrated that Polo-like kinase 1 (PLK1) activity facilitates CENP-A assembly during interphase in human cells by promoting the complete assembly of the CENP-A chaperone complex at centromere (*9*, *11*, *25*). Therefore, we tested whether PLK1 is essential for centromere architecture. In interphase egg extracts, inhibition of PLK1 with BI-2536 after exit from M phase decreased the total levels of CENP-A by nearly 50%, similar to results observed in human cell culture (Figs. 2A and 2B). To confirm that PLK1 inhibition blocks new CENP-A assembly in the *Xenopus* egg extract system, we expressed 3X-Myc-CENP-A and HJURP from mRNA and analyzed 3X-Myc-CENP-A incorporation at centromeres after entry into interphase (Figs. S3A-S3D). In the DMSO control, 3X-Myc-CENP-A was incorporated at centromeres; in BI-2536-treated cells, its incorporation was blocked (Fig. S3A and S3B). Furthermore, BI-2536 treatment blocked Mis18α localization α to centromeres (Fig. S3C and S3D), highlighting the conservation of PLK1-dependent CENP-A assembly. Strikingly, PLK1 inhibition completely abrogated the formation of CENP-A, CENP-C, and CENP-T rings (Figs. 2C and 2D). Notably, the levels of CENP-T and CENP-C at centromeres were not affected by treatment with BI-2536 (Figs. S3E and S3F), which agrees with previous findings that a full complement of CENP-A is not required for CCAN assembly (*17*, *26*, *27*). Thus, our data demonstrate that PLK1-dependent control of the CENP-A chaperone complex is not required for CCAN assembly, but raises the possibility that it is necessary for the spatial arrangement of the centromere and the CCAN.

### HJURP controls centromere architecture independently of its role in CENP-A assembly

PLK1 inhibition prevents CENP-A assembly by blocking the centromere localization of the Mis18 complex and HJURP during interphase (*9*, *11*, *25*). To directly test whether the chaperone complex is required for the spatial arrangement of centromeres, we immunodepleted HJURP from egg extracts (Fig. S4A). As expected (*7*, *28*), HJURP depletion prevented CENP-A assembly, as indicated by a nearly 50% decrease in total CENP-A at centromeres (Figs. S4B and S4C). HJURP depletion severely decreased the frequency of CENP-T ring formation in interphase, with CENP-T instead forming punctate structures (Fig. 3A and Fig. S4D). Furthermore, the decrease of chromatin density in the central cavity of the centromere was lost upon HJURP depletion (Figs. S4E-G). Thus, HJURP is required for the spatial arrangement of the centromere and for the clearing of chromatin at centromeres in interphase.

**Fig. 3.**
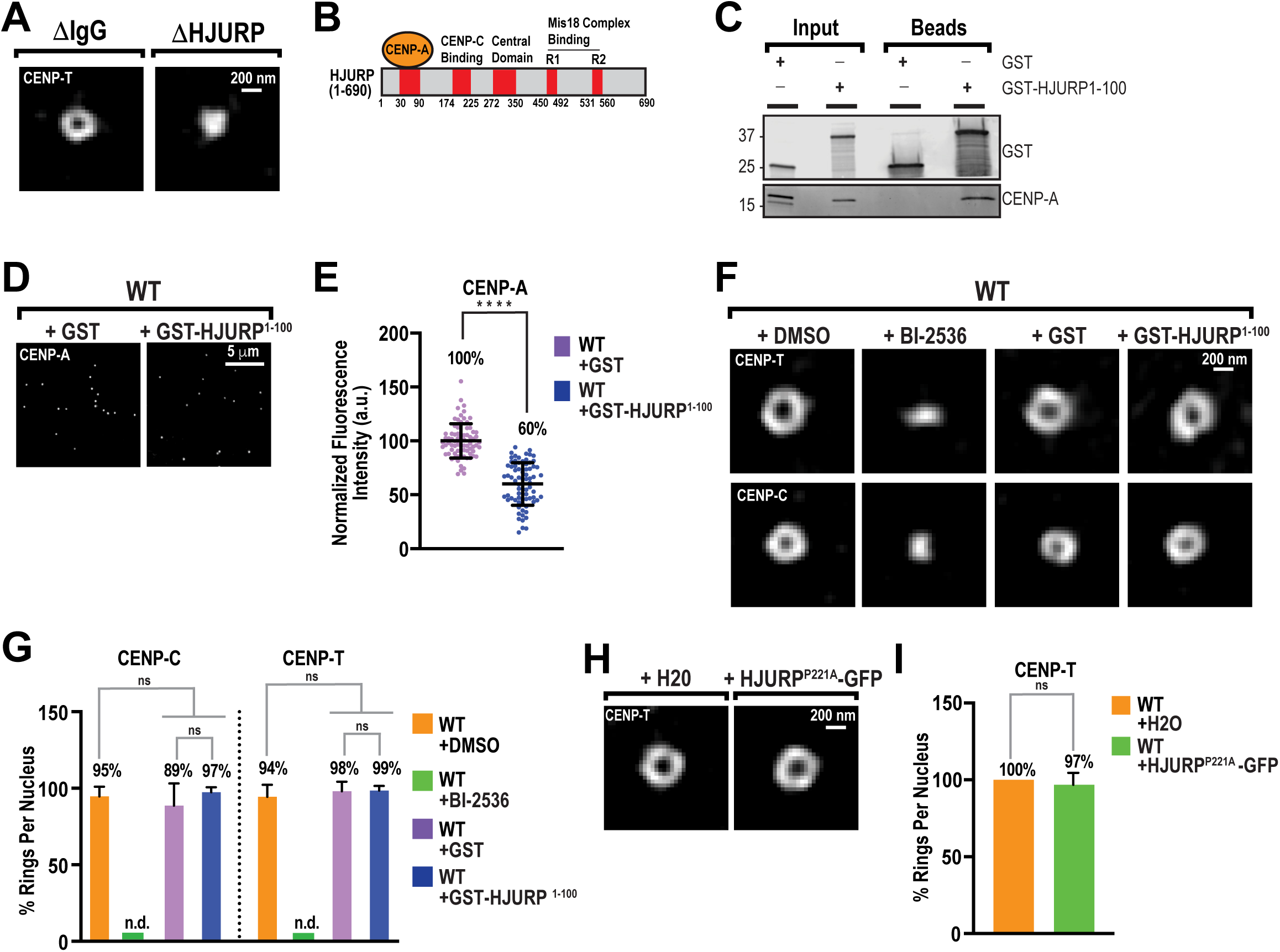
HJURP is required for centromere organization independently of CENP-A assembly. (**A**) Representative 3D-SIM images of CENP-T on interphase nuclei, at *t*=90’, in mock depleted (ΔIgG) and HJURP depleted (ΔHJURP) extracts. Scale bar is 200 nm. (**B**) Domain organization of HJURP showing CENP-A/H4 dimer, CENP-C, and Mis18 complex binding regions and the central domain. (**C**) Western blot for proteins that co-purify with GST- or GST-HJURP^1-100^-coupled glutathione beads from high-speed supernatant egg extracts in the absence of chromosomes. Input is 5% of the total extract used per co-purification. (**D**) Representative immunofluorescence images of CENP-A on interphase nuclei, at *t*=90, in extracts with purified GST or GST-HJURP^1-100^ added. Scale bar is 5 µm. (**E**) Quantification of fluorescence intensity of CENP-A shown in (D) normalized to GST. n = 75 centromeres per condition. A.U., arbitrary units. Error bars represent SD, ****, *P* < 0.0001. (**F**) Representative 3D-SIM images of CENP-C and CENP-T on interphase nuclei, at *t*=90’, in extracts with DMSO, BI-2536, GST, and GST-HJURP^1-100^. Scale bar is 200 nm. (**G**) Percentage of observable ring-like structures per nucleus for conditions in (F). n=10 nuclei per condition, and nβ145 centromeres total. Error bars represent SD. NS = not significant. (**H**) Representative 3D-SIM images of CENP-T on interphase nuclei, at t=90’, in extracts with either water (H2O) or HJURP^P221A^-GFP mRNA added. (**I**) Percentage of observable ring-like structures per nucleus for conditions in (H). n=10 nuclei per condition, and nβ145 centromeres total. Error bars represent SD. NS = not significant.

We next sought to test whether new CENP-A assembly is required for centromere and CCAN organization. To test this, we sought to perturb HJURP’s ability to recruit CENP-A/histone H4 heterodimers without affecting its localization to centromeres (*29*). We purified the CENP-A/histone H4-binding Scm3 domain of HJURP fused to GST (Fig. 3B; GST-HJURP^1-100^) from bacteria and added it to extracts at a final concentration of 5 μM. Since HJURP is stored in a complex with soluble CENP-A/H4 in egg extracts at low levels (*30*), this concentration should be in excess of endogenous HJURP. It therefore should sequester CENP-A/H4 heterodimers away from endogenous HJURP. Indeed, we found that GST-HJURP^1-100^, but not GST, effectively pulled down soluble CENP-A from extracts in the absence of nuclei (Fig. 3C). Addition of GST-HJURP^1-100^ to interphase extracts reduced total CENP-A at centromeres to levels similar to those observed after PLK1 inhibition (Figs. 3D and 3E). However, unlike PLK1 inhibition, GST-HJURP^1-100^ did not perturb the localization of Mis18α-GFP to centromeres (Fig. S4H and S4I). Thus, GST-HJURP^1-100^ acts in a dominant-negative manner, blocking CENP-A assembly without perturbing the recruitment of chaperone complexes to centromeres. Remarkably, GST-HJURP^1-100^ did not affect the formation of CENP-C or CENP-T rings, whereas PLK1 inhibition completely blocked ring formation (Figs. 3F and 3G). To test whether CENP-A assembly can modulate the CCAN architecture, we sought to increase the amount of newly assembled CENP-A. Prior work has shown that the HJURP^P221A^ mutation significantly increases new CENP-A assembly at centromeres (*31*). In agreement with these findings, expression of the hyperactive HJURP^P221A^ mutant in interphase extracts led to a 64% increase in CENP-A levels at centromeres over wild-type (Figs. S4J and S4K). However, HJURP^P221A^-GFP did not affect the formation of CENP-T rings (Figs. 3H and 3I). Altogether, our data suggest that the chaperone complex, but not its function in CENP-A assembly, is required for the 3D organization of centromeres and the CCAN.

### The CENP-A chaperone complex resides within the central cavity of centromeric chromatin structures

Current models of CENP-A assembly propose that HJURP and the Mis18 complex localize to pre-existing CENP-A nucleosomes, replacing nearby histone H3.3 “placeholder” nucleosomes with new CENP-A molecules from the soluble pool (*13*, *32–36*). This model implies that these chaperone complexes should localize proximal to the CCAN and CENP-A and have a similar spatial arrangement. To test this, we used 3D-SIM imaging combined with anti-GFP nanobody labeling to analyze the spatial distribution of GFP-tagged chaperone complex proteins relative to the CCAN. Unlike CENP-A and the CCAN, which form continuous ring-like structures in the XY plane, each member of the chaperone complex is localized as discrete punctate structures within the interior of CENP-T rings (Fig. 4A). Line-scan analysis showed a unimodal distribution of fluorescence intensity for both HJURP-GFP and Mis18BP1-GFP at centromeres, peaking at the local minimum between the bimodal peaks observed for CENP-T fluorescence (Figs. 4B and 4C). In contrast to the chaperone machinery, CENP-T-GFP formed ring-like structures similar to those observed with antibodies targeting endogenous CENP-T and CENP-C, demonstrating that anti-GFP nanobody labeling does not interfere with the ability to observe such structures (Fig. 1B and Figs. S5A-C). By metaphase, these CENP-T rings collapsed, and HJURP fluorescence signals were no longer detectable at centromeres (*31*) (Figs. S5D and S5E). Together, our data indicate that the HJURP chaperone complex occupies the central cavity enclosed by CENP-A and CCAN structures during interphase, and the disappearance of these chaperone proteins at mitosis coincides with the structural collapse of this cavity.

**Fig. 4.**
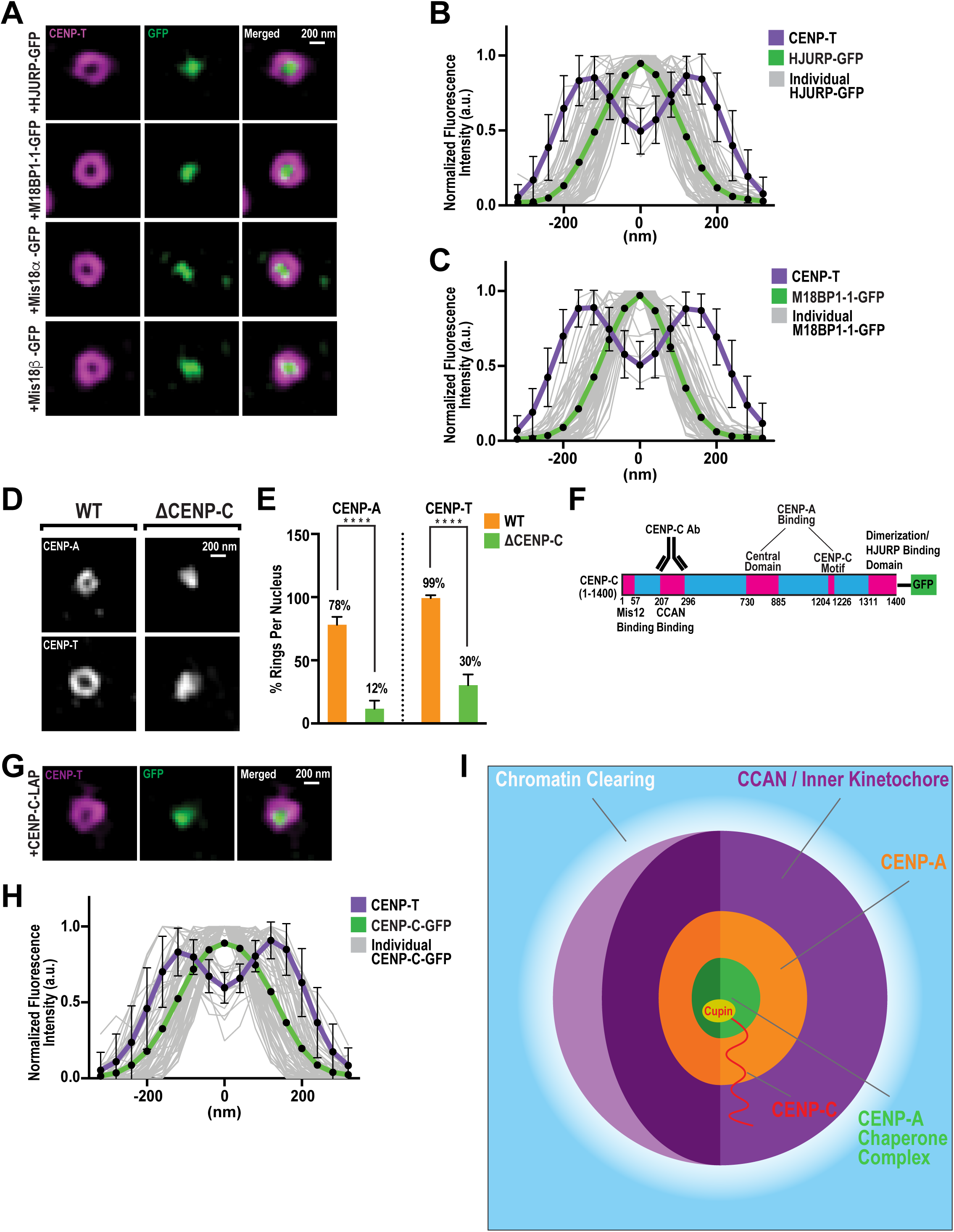
CENP-C and the CENP-A chaperone complex spatially organize centromeric chromatin. (**A**) Representative 3D-SIM images of CENP-T (magenta) and GFP (green, GFP-Booster ATTO 488) on interphase nuclei, at t=90’, assembled from extracts containing indicated *in vitro* translated GFP-tagged chaperone complex proteins. Scale bar is 200 nm. (**B**) Linescan quantification of CENP-T and HJURP-GFP fluorescence at centromeres from (A). Bold lines indicate average intensity, and thin gray lines indicate individual traces of HJURP-GFP fluorescence. Each trace was aligned by the central minimum and normalized to the maximum intensity. n=75 kinetochores. Error bars represent SD. (**C**) Linescan quantification of CENP-T and M18BP1-1-GFP fluorescence at centromeres from (A), analyzed as in (B). n=75 Kinetochores. Error bars represent SD. (**D**) Representative 3D-SIM images of CENP-A and CENP-T on interphase nuclei, at *t*=90’, assembled from WT or CENP-C depleted (ΔCENP-C) extracts. Scale bar is 200 nm. (**E**) Percentage of observable ring-like structures per nucleus for conditions in (F). n=10 nuclei per condition, and n≥145 centromeres total. Error bars represent SD, ****, *P* < 0.0001. (**F**) Domain organization of CENP-C showing Mis12-binding, CCAN-binding, central domain, CENP-C motif, HJURP-binding and dimerization domains. Antibody epitope binds to aa 207-296, as indicated. (**G**) Representative 3D-SIM image of CENP-T (magenta) and GFP (green) on interphase nuclei, at *t*=90’, in extracts containing *in vitro* translated CENP-C-GFP. Scale bar is 200 nm. (**H**) Linescan quantification of CENP-T and CENP-C-GFP fluorescence at centromeres from (G), analyzed as in (B). n=75 Kinetochores. Error bars represent SD. (**I**) Schematic of centromere and kinetochore organization during interphase. Concentric shells of CENP-A-enriched chromatin and CCAN define a region of decreased chromatin density. The CENP-A chaperone machinery occupies the central cavity where chromatin is most decreased. CENP-C spans the entire structure, with its C-terminal cupin domain inside the central cavity, and its N-terminal region within the CENP-A/CCAN shells.

### CENP-C forms a molecular scaffold that connects CENP-A nucleosomes to chaperone complexes to promote centromere organization

CENP-C directly connects pre-existing CENP-A chromatin with the CENP-A chaperone complex (*7*, *31*, *37–39*), suggesting it plays a role in centromere and kinetochore architecture during interphase. To test this suggestion, we depleted CENP-C from egg extracts. As expected, CENP-C depletion prevented new CENP-A assembly, reducing total centromeric CENP-A levels by nearly 50% compared to controls (Fig. S6A-C). Remarkably, loss of CENP-C drastically decreased the frequency of CENP-A and CENP-T rings (Figs. 4D and 4E). Instead, in the absence of CENP-C, CENP-A and CENP-T adopted irregular punctate distributions (Fig. 4D), resembling the early interphase spatial arrangement of CENP-T and CENP-C (Fig. 1A). In contrast, depletion of CENP-T minimally affected the formation of CENP-A and CENP-C rings (Figs. S6D-S5H). These findings demonstrate that CENP-C specifically controls the higher-order, three-dimensional organization of CENP-A and the CCAN during interphase.

Given the direct interaction between CENP-C and HJURP (*31*, *40*),it was surprising that HJURP exhibited a strikingly different localization pattern compared to CENP-C (Figs. 1B and 4A). CENP-C is a modular scaffold protein containing distinct, evolutionarily conserved domains that bind the outer kinetochore, the CCAN, CENP-A nucleosomes, and HJURP (*31*, *41*, *42*)(Fig. 4F). Our anti-CENP-C antibody targets an epitope within CENP-C that overlaps with the CENP-HIKM and CENP-NL binding regions (*42*, *43*). This epitope is approximately 1,000 residues away from its C-terminal cupin domain, which is responsible for HJURP binding (*31*), suggesting that the CENP-C C-terminus may localize differently from the rest of the CCAN. To test this, we tagged the CENP-C C-terminus with GFP immediately downstream of the cupin domain and determined its localization relative to the CCAN using an anti-GFP nanobody. Strikingly, the CENP-C C-terminus localized within the center of CENP-T rings, exhibiting a unimodal fluorescence distribution similar to that observed for HJURP-GFP and M18BP1-GFP (Figs. 4G and 4H). Thus, CENP-C adopts an extended and likely radially oriented conformation, positioning its HJURP-binding C-terminus within the central cavity of the centromeric chromatin structure. At the same time, its CCAN-scaffolding N-terminus forms a peripheral shell around CENP-A chromatin. Collectively, our findings reveal a previously unrecognized role for CENP-C and the chaperone complex in organizing the architecture of centromeres and kinetochores in interphase and suggest that this process is important for their mitotic organization.

## Discussion

Advances in assembling repetitive sequences have clarified where CENP-A resides along centromeric DNA (*44*, *45*), but how these one-dimensional positions are organized into a three-dimensional, functional centromere is less well understood (*46*). We reveal a structural role for the CENP-A chaperone complex that is independent of its canonical deposition function. In interphase, HJURP and the Mis18 complex occupy a chromatin-poor central cavity encircled by concentric shells of CENP-A nucleosomes and the CCAN. The CCAN protein CENP-C spans this assembly in a defined N-to-C orientation, linking the chaperone core to CENP-A chromatin and the broader CCAN, and is required for formation of interphase centromere architecture (Fig. 4I).

Our data indicate that a cell-cycle-dependent mechanism regulates centromere architecture. PLK1-dependent chaperone assembly at centromeres during interphase creates a chromatin clearing encircled by CENP-A-enriched chromatin and the CCAN. Prior work shows that PLK1-driven assembly is antagonized by CDK, which evicts HJURP and other chaperone components via phosphorylation (*5*, *9*, *11*, *13*, *25*). In *Xenopus* egg extracts, chaperone eviction occurs upon mitotic entry, driven in part by CDK-dependent disruption of the HJURP-CENP-C interaction (*31*). Consistent with this, we observe that loss of the shell-like architecture in mitosis coincides with loss of HJURP from centromeres (Fig. S5D), supporting a model in which CDK-dependent chaperone removal in mitosis triggers the collapse of the extended CENP-A/CCAN shells into smaller puncta that serve as sites for outer kinetochore assembly. In somatic human cells, where CENP-A loading is confined to G1, CDK activity during S and G2 is sufficient to reverse PLK1-enabled recruitment (*12*), predicting the collapse of centromere structure during S/G2. Indeed, super-resolution imaging in U2OS cells revealed rosette-like CENP-A organization surrounding HJURP in G1, which was lost after G1 exit (*21*). Thus, our findings suggest that PLK1-dependent formation of centromere shells and their CDK-dependent collapse are general features of vertebrate centromeres.

The interphase organization of centromeric chromatin and the CCAN likely impacts centromere and kinetochore function in mitosis. Studies in human cells have defined mitotic centromere architecture: one showed that individual centromeres comprise two subdomains of chromatin loops forming a bipartite structure stabilized by the SMC complexes condensin and cohesin (*19*); another identified chromatin clearings that demarcate CENP-A and inner kinetochore (CCAN) territories and demonstrated a requirement for CENP-C to maintain these clearings (*20*). Complementary work reported ring-like arrangements of α-satellite DNA and CENP-A at mitotic centromeres in human cells (*47*), and that outer kinetochore proteins form ring-like structures in mitotic *Xenopus* egg extracts (*48*). Against this backdrop, our results support a model in which the chaperone complex creates chromatin clearings surrounded by radially organized shells of CENP-A and the CCAN in interphase, and these clearings function as “placeholders” into which CENP-A chromatin and the CCAN redistribute after chaperone eviction, generating the mitotic chromatin clearings and architectures described by others. This chaperone-dependent architecture formed in interphase may also template assembly of the outer kinetochore, thereby promoting proper attachment of kinetochores to spindle microtubules. We emphasize that our evidence establishes temporal correlation and architectural continuity but does not directly demonstrate that interphase clearings are the immediate precursors of mitotic clearings.

The interphase arrangement of CENP-A and the chaperone complex may also be necessary for efficient and accurate CENP-A assembly. A predominant model posits that a pre-existing CENP-A nucleosome templates assembly of a new CENP-A nucleosome to restore pre-replicative CENP-A levels; proximal histone H3.3 nucleosomes serve as “placeholders” that are evicted and replaced by HJURP-mediated delivery of CENP-A/H4 heterodimers (*13*, *32–36*). How this eviction and replacement are targeted, and how centromere drift and expansion are prevented, remain unclear (*46*). Although we demonstrate that CENP-A assembly is not required for centromere architecture, the shell-like architecture may regulate assembly in two ways. First, the high local concentration of chaperone complexes in a defined space near pre-existing CENP-A nucleosomes may create “factories” that accelerate eviction and assembly. Second, sequestering the chaperone complex away from bulk chromatin could prevent loading of new CENP-A at non-centromeric sites and excessive loading at centromeres. While heterochromatin and other factors act as local boundaries to prevent centromere drift and expansion (*49*, *50*), current models emphasize one-dimensional spreading along linear DNA. Our results raise the possibility that centromere drift and expansion are further delimited in three dimensions by the spatial organization of CENP-A chromatin and chaperone complexes, potentially creating a three-dimensional zone of eviction and assembly.

How might the chaperone complex and CENP-C organize centromeric chromatin into shell-like structures around chromatin clearings in interphase, independent of new CENP-A assembly? Our findings suggest that the chaperone complex and CENP-C exclude bulk chromatin while maintaining the proximal organization of CENP-A-enriched chromatin. Phase separation and electrostatic repulsion have been proposed to promote centromere organization through self-association of centromeric chromatin and repulsive forces generated by kinetochore proteins (*51*). Additionally, the cupin domain of CENP-C, which interacts with HJURP (*31*, *52*), forms higher-order oligomers that promote self-association of centromeric chromatin (*53*). It is therefore possible that CENP-C oligomers cluster CENP-A nucleosomes and that chaperone complexes radially repel these clusters to form shells. However, the oriented N-to-C arrangement of CENP-C molecules we observe (Figs. G-I) and the defined stoichiometry of the HJURP/Mis18 complex (*33*, *54*) argue against liquid-liquid phase separation, which relies on largely non-specific, dynamic interactions, as the main organizing principle (*55*). Alternatively, the chaperone complex may form a rigid structural foundation tethered to centromeric chromatin through CENP-C. This model is analogous to centriole organization (*56–58*), in which proteins such as SAS-6 provide a defined nine-fold symmetric architecture to organize centriolar microtubules, and pericentrin serves as a flexible yet oriented scaffold that connects the centriole wall to the pericentriolar material. Thus, centromeres and kinetochores may share similar principles of organizational control with centrioles and centrosomes.

## Acknowledgments

We thank all members of the Akera, Kelly, Rosin, and Rusan labs for helpful scientific discussions. We thank A. Losada, A. Straight, A. Arnaoutov, and M. Dasso for kindly providing reagents. We thank T. Akera for critical reading of the manuscript. We thank X. Wu and the National Heart, Lung, and Blood Institute light microscopy core facility for their continuous support with 3D-SIM imaging on the DeltaVision OMX SR microscope. We thank M. Kruhlak and the light microscopy core facility of CCR at the National Cancer Institute for their technical assistance in super-resolution microscopy (Zeiss Elyra LSM 780, Zeiss LSM 880 Airyscan, and Nikon SoRa Spinning Disk) imaging. The deep learning work was performed and supported by the Advanced Imaging and Microscopy Resource of the National Institute of Biomedical Imaging and Bioengineering. This work was supported by the Howard Hughes Medical Institute (HHMI). This article is subject to HHMI’s Open Access to Publications policy. HHMI laboratory heads have previously granted a non-exclusive CC BY 4.0 license to the public and a sub-licensable license to HHMI in their research articles. Pursuant to those licenses, the author-accepted manuscript of this article can be made freely available under a CC BY 4.0 license immediately upon publication.

## Funding

Division of Intramural Research at the National Institutes of Health, the National Cancer Institute, Center for Cancer Research, ZIA BC011491 (A.E.K.)

Howard Hughes Medical Institute (H.S.)

## Author contributions

Conceptualization: H.G., A.E.K.

Methodology: H.G., H.S., A.E.K.

Investigation: H.G., J.H., Y.W., J.C., G.K.

Supervision: H.S., A.E.K.

Visualization: H.G., J.H.

Writing – original draft: H.G., A.E.K.

Writing – review & editing: H.G., J.H., Y.W., J.C., H.S., A.E.K.

## Data and materials availability

All data are available in the main text or supplementary materials. Materials generated in the current study are available from the corresponding author on reasonable request.

## Materials and Methods

### Cloning and plasmid construction

*Xenopous laevis* M18BP1 (isoform M18BP1.L), Mis18ɑ, and Mis18β genes were synthesized (Twist Bioscience) and inserted into a pCS2+ expression vector containing a LAP tag, a combination of 6xHis tag and EGFP, at the C-terminal. 3xMyc-CENP-A was synthesized and inserted into a pCS2+ expression vector (Twist Bioscience). CENP-T, HJURP and CENP-C were PCR amplified and cloned into a pCS2+ vector containing a C-terminal LAP tag by Gibson assembly (NEB). Residues 1-100 of HJURP (HJURP^1-100^) were synthesized and inserted (Twist Bioscience) into the pGEX-6P-1 vector. HJURP mutation P221A (*31*) was generated by Q5 site-directed mutagenesis (NEB) using the primer 5’-CAAAGTCTCTGCCATGAAATTTCGAAG - 3’.

### Generation of anti-xCENP-A serum for Western Blotting

To generate serum against xCENP-A for Western Blotting, a peptide spanning residues 1 to 20 of *Xenopus laevis* CENP-A including an additional cysteine residue for conjugation was synthesized, MRPGSTPPSRRKSRPPRRVSC-amide (Biosynth, MA, USA), and used to immunize rabbits (Labcorp Early Development Laboratories Inc., WI, USA).

### Assembly of interphase nuclei and mitotic chromosomes in *Xenopus* egg extracts

Metaphase-arrested *Xenopus laevis* egg extract was prepared as previously described (*59*). To examine *de novo* kinetochore assembly, *Xenopus laevis* sperm nuclei and CaCl_2_ were added to the egg extract to release it into interphase, and incubated for 90 minutes at 21°C. Reactions were then driven into M phase by dilution with two volumes of CSF-arrested extract and incubated for another 40-45 minutes at 21°C. Interphase nuclei or mitotic chromosomes were collected at various time points as indicated, and processed for immunofluorescence staining.

### Immunodepletion of *Xenopus* egg extracts

Immunodepletions were performed using antibodies bound to Protein A beads (Dynabeads; Invitrogen). For immunodepletion of CENP-C or HJURP, 100 µl of extract was incubated for one or two rounds with 30 µg of anti-CENP-C or anti-HJURP antibody (*28*, *60*) (a gift of Ana Losada, CNIO) bound to 100 µl of beads on ice for 45 minutes. For immunodepletion of CENP-T, 100 µl of extract was incubated for two rounds with 15 µg anti-CENP-T antibody (*60*) (a gift of Ana Losada, CNIO) bound to 100 µl of beads on ice for 45 minutes. For mock depletions, 100 µl of extract was incubated for two rounds with 10 µg of IgG (Sigma, I5006).

### Expression of exogenous mRNA in egg extracts

3xMyc-CENP-A and HJURP mRNAs were transcribed in vitro by using the mMessage Machine SP6 kit (Thermo Fisher Scientific). To prevent the translation of endogenous RNAs, RNase A (Ambion) was added to the extract at a final concentration of 100 ng/ml for 15 minutes at 12°C. An RNase inhibitor (RNasin Plus RNase Inhibitor; Promega) was added at a 1:50 dilution for 5 minutes at 12°C. Yeast tRNAs (50 μg/ml) were added, and the extract was kept on ice.

### DNA replication assay in *Xenopus* egg extracts

To study DNA replication, DMSO or 100 µg/ml aphidicolin (Sigma, A0781) were added to extracts supplemented with 10 µM Cy3-dUTP (Cytiva, PA53022). Then, sperm nuclei and CaCl_2_ were added to the reactions and incubated at 21°C for 90 minutes, and then processed for fluorescence imaging.

### CENP-A loading assay in *Xenopus* egg extracts

To monitor the assembly of new CENP-A molecules at centromeres, 500 ng of 3xMyc-CENP-A and 500 ng of HJURP mRNAs were added per 20 µl of CSF-arrested extract and incubated at 21°C for 30 minutes to translate the 3xMyc-CENP-A and HJURP, as described previously (*7*). Cycloheximide was then added to the reactions to block further protein translation, followed by the addition of sperm nuclei and CaCl_2_ to release into interphase. The reactions were then incubated for 90 minutes at 21°C. Reactions were then processed for immunofluorescence. For experiments to examine the effects of PLK1 inhibition on CENP-A loading, DMSO or 10 µM BI-2536 (Selleckchem, S1109) were added to the indicated reactions 30 minutes after the addition of CaCl_2_ to allow for exit from M phase (*61*).

### Immunofluoresence and Widefield Microscopy

To immunostain kinetochores, *Xenopus* egg extract was fixed for 5 min by 20-fold dilution in BRB80, 20% glycerol, 0.5% Triton X-100, and 3.7% formaldehyde at room temperature. Fixed reactions were layered onto a cushion of BRB80 and 40% glycerol overlaying poly-L-lysine– coated coverslips (No. 1) placed in a 24-well plate. Nuclei were adhered onto coverslips on plate holders at 4,000 rpm for 15 min at 18°C in a centrifuge (Eppendorf 5810R). Cushions were washed with BRB80. For immunostaining of CENP-A, CENP-C, and CENP-O, coverslips were postfixed in ice-cold methanol for 5 min, blocked with Abdil (TBS, 0.1% Tween20, 2% BSA, and 0.1% sodium azide) overnight at 4°C then incubated in the primary antibody at room temperature for 1 h followed by secondary antibody (Alexa Fluor 488) at room temperature for 1 h unless otherwise noted. For any immunostaining involving CENP-T, no methanol fixation was used, and Abdil with 3% BSA was used. For co-immunostaining of CENP-T or CENP-C and Myc- or LAP-tagged proteins, coverslips were blocked with Abdil with 3% BSA at room temperature for 1.5 h, then incubated with Myc primary antibody or GFP-Booster Atto 488 at 18°C overnight, then incubated with secondary antibody for 1h at room temperature. Coverslips were then incubated successively with CENP-T or CENP-C primary and secondary antibodies for 1 h at room temperature. All washes and antibody dilutions were done with AbDil buffer. Nuclei were stained with Hoechst 33258 unless otherwise noted. Coverslips were mounted on a glass slide in 80% glycerol in a PBS medium. The following antibodies were used at the indicated dilutions: CENP-C (*60*) (a gift of Ana Losada, CNIO), 1:150; CENP-T (*60*) (a gift of Ana Losada, CNIO), 1:100; CENP-A (*62*) (a gift of Aaron Straight, Stanford University), 1:500; CENP-O (a gift of Alexei Arnoutov and Mary Dasso, NICHD, NIH), 1:200; anti-Myc (Santa Cruz, sc-40), 1:100; GFP-Booster Atto 488 (Chromotek, gba488) 1:400; Alexa Fluor 488 (Jackson Immuno Research, 711-545-152), 1:400; Alexa Fluor 647 (Jackson Immuno Research, 711-605-152), 1:400. For quantification of immunofluorescence, samples were imaged with a 0.2 or 0.4 µm step size using an Eclipse Ti (Nikon) composed of a Nikon Plan Apo ×100/ 1.45, oil immersion objective, a Plan Apo 40×/0.95 objective, and a Hamamatsu Orca-Flash 4.0 camera. Images were captured and processed using Nikon’s NIS Elements AR 4.20.02 software and analyzed in ImageJ/Fiji. The acquired Z-sections were converted into a maximum intensity projection using NIS Elements and Fiji. Kinetochore intensity was measured using Fiji centering 9 × 9- and 13 × 13-pixel regions over individual kinetochores. Total fluorescence intensity was recorded from each region. To correct the background fluorescence, the difference in intensities between the two regions was determined and then made proportionate to the smaller region. This background value was then subtracted from the smaller region to determine kinetochore intensity with background correction, as previously reported (*63*).

### Super-resolution microscopy data acquisition and reconstruction

Unless otherwise noted, 3D-Structured illumination microscopy (3D-SIM) imaging was performed on a GE DeltaVision OMX SR microscope (GE Healthcare) equipped with a 60x 1.42 NA oil objective (Olympus) using DeltaVision immersion oil with a refractive index of 1.514. Z-stacks were acquired for a single channel at 0.125 µm increments. Raw data were reconstructed using SoftWorx software (GE Healthcare) with a Wiener filter constant set to 0.003. Lasers with 405, 488 nm, and 640 nm wavelengths were used to image DAPI, Alexa 488, or GFP-Booster Atto 488 and Alexa 647.

Structured illumination (SIM) super-resolution images (Fig. S1E, left panel) were acquired using a Zeiss LSM780 Elyra SIM microscope equipped with a 63x plan-apochromat (NA 1.4) oil immersion objective lens. Images were collected with a 0.032 µm x-y pixel size, z-step size of 0.090 µm, 5 phases, and 3 rotations of the SIM illumination grating. Images were processed using the SIM image processing algorithm in the Zeiss Zen software (v.13.0). A 488 nm laser was used to image Alexa Fluor 488.

Airyscan super-resolution images (Fig. S1E, right panel) were acquired using a Zeiss LSM880 microscope equipped with a 63x plan-apochromat (NA 1.4) oil immersion objective lens. Images were collected with a 0.044 µm x-y pixel size and 0.160 µm z-step size. The images were processed using the Airyscan processing algorithm in the Zeiss Zen software (v. 14.0.18). A 488 nm laser was used to image Alexa Fluor 488.

Nikon SoRa super-resolution images (Fig. S1E, middle panel) were acquired using a Nikon SoRa spinning disk microscope equipped with a 60x Apo TIRF (NA 1.49) oil immersion objective lens and Photometrics BSI sCMOS camera. Images were collected with a 0.03 µm x-y pixel size and 0.15 µm z-step size and deconvolved using a modified Lucy-Richardson constrained iterative deconvolution algorithm in the Nikon Elements image analysis software (v. 5.4.1). A 488 nm laser was used to image Alexa Fluor 488.

### 3D-SIM image analysis

ImageJ/Fiji was used for the 3D-SIM image analysis. A single plane from a z-stack of a single or double channel image stack was used for image analysis and visualization. Unless otherwise noted, a representative image of a single kinetochore was cropped and shown in the figures. An orthogonal view (XZ or YZ) of an XY plane was created using Fiji’s orthogonal views tool. The line intensity profile across the kinetochore structure was measured using Fiji’s plot profile tool. The mean background intensity for centromeres and kinetochores was measured from the region within the nucleus that lacked a centromere or kinetochore, whereas for DAPI, it was estimated from the exterior of the nucleus. The intensity profile of a single channel was generated by subtracting the mean background intensity per pixel from the total intensity per pixel along a line intensity profile. The background-corrected intensity profile was then normalized to the maximum intensity, and the minimum or maximum peak of the intensity profile was recorded as a center of the curve and plotted against the distance from the center. The maximum width of a kinetochore ring was defined by measuring the distance between the two highest peaks in the intensity profile generated from a line scan across the longest axis of the ring structure.

### Deep learning analysis of 3D-SIM data

Using neural networks to improve axial resolution was described previously (*24*). Model training and model application were performed on an NVIDIA RTX A6000 GPU (with 48 GB memory) installed on a local workstation. We used Python version 3.9.18 for all neural networks. For training, we first interpolated the 3D-SIM volumes to 40-nm pixel size in all three dimensions. The interpolated volumes were blurred with a 1D Gaussian function, sigma = 2.4 pixels along the x dimension. Next the interpolated volumes were blurred with a 1D Gaussian function (sigma = 1.0 pixel) along the z dimension to remove spurious sidelobes in the Fourier domain. The purpose of these blurring operations is to degrade lateral resolution so it resembles the native axial resolution of the 3D SIM system. Stacks were then down sampled along the x dimension to mimic the coarse axial sampling of the experiment; and up-sampled to recover an isotropic pixel size, serving as the ‘input’ for training for the CARE network (*64*). The goal of the network is to reverse this image degradation, thereby improving resolution along the degraded x direction. Patches of size 64 × 64 × 64 were randomly cropped from the data, and 15% of the patches were set aside for validation. The training learning rate was 2 × 10^4^; the number of epochs was 100; the number of steps per epoch was 200; and the mean absolute error (MAE) was used as loss function. To apply the trained model, we (1) interpolated, blurred, downsampled, and upsampled 3D-SIM volumes as described above; (2) rotated the resulting volumes at six distinct angles about the ‘y’ axis; (3) applied the network to each rotated volume to increase resolution in 1D; (4) rotated the network predictions back to their original orientation; (5) cropped them to their original size; (6) Fourier transformed each volume; (7) computed the maximum value at each spatial frequency across all rotations; and (8) obtained the final result by applying inverse Fourier transform and taking the absolute value.

### *In vitro* transcription and translation

The SP6-coupled *in vitro* transcription and translation (TNT) system (Promega) was used to express CENP-T-LAP, CENP-C-LAP, HJURP-LAP, Mis18ɑ-LAP, Mis18β-LAP, and M18BP1-LAP from 2 µg of plasmid DNA. To assess the localization of LAP-tagged protein, 2 μl of the TNT reaction was added to 20 μl of the indicated extract condition at the beginning of interphase.

### Purification of GST-HJURP^1-100^ and GST

GST-HJURP^1-100^ or GST was expressed in BL21 (DE3) Rosetta-2 cells. Expression was induced with 0.5 mM IPTG for 2.5 h at 37°C. Lysates containing GST-HJURP^1-100^ or GST were bound to Glutathione Sepharose 4B resin and eluted with 20 mM HEPES, 20 mM reduced Glutathione, pH 7.9, 500 mM NaCl, 1 mM TCEP, and exchanged into XB buffer (0.1 M KCl, 0.1 mM MgCl_2_, 50 mM Sucrose and 10 mM HEPES, pH 7.7, 1 mM TCEP). GST-HJURP^1-100^ or GST were added to the extract at a final concentration of 5 µM.

### Co-immunoprecipitations

Immunoprecipitations were performed to determine the binding of GST-HJURP^1-100^ to soluble CENP-A in high-speed supernatant (HSS) *Xenopus* egg extracts, which were prepared as described previously (*65*). Purified GST-HJURP^1-100^ or GST were added to 50 µl of HSS extracts without chromatin to a final concentration of 5 µM, and incubated at 21°C for 30 minutes. 50 µl of Glutathione Sepharose 4B Beads were washed four times in binding buffer (20 mM HEPES, pH 7.7, 150 mM NaCl, 1mM TCEP) and added to the extracts, and then incubated with end-over-end rotation for 1h at 4°C. Beads were briefly spun and washed four times with wash buffer (20 mM HEPES, pH 7.7, 150 mM NaCl, 1 mM TCEP, 0.01% NP40) before eluting with 2X sample buffer.

### Western Blots

Primary antibodies were diluted in Licor blocking solution with a final Tween-20 concentration of 0.1%. The following antibodies and antibody dilutions were used: anti-CENP-T (a gift of Ana Losada, CNIO), 1:250; anti-CENP-C (a gift of Ana Losada, CNIO) 1:250; anti-HJURP (a gift of Ana Losada, CNIO) 1:200; GST (Pierce, 700775) 1:1000; anti-CENP-A serum (this paper) 1:100; anti-histone H3T3ph (Epitomics, 2162–1) 1:10,000; anti-Myc (Santa Cruz, sc-40), 1:00; anti-α-tubulin (DM1, 1:20,000; Sigma). Secondary antibodies from Licor were used (goat anti-rabbit 800 nm and goat anti-mouse 680 nm), as was the Licor imaging system to scan membranes.

### Quantification and statistical calculation

All graphs were prepared in GraphPad Prism 10 Software. Data are presented as mean ± s.d. Statistical analysis between two data sets was performed using the unpaired t-test of Prism. Sample size (n=), statistical parameters, and p values are provided in the figures and legends.

## Supplementary Figure Legends

**Fig. S1.**
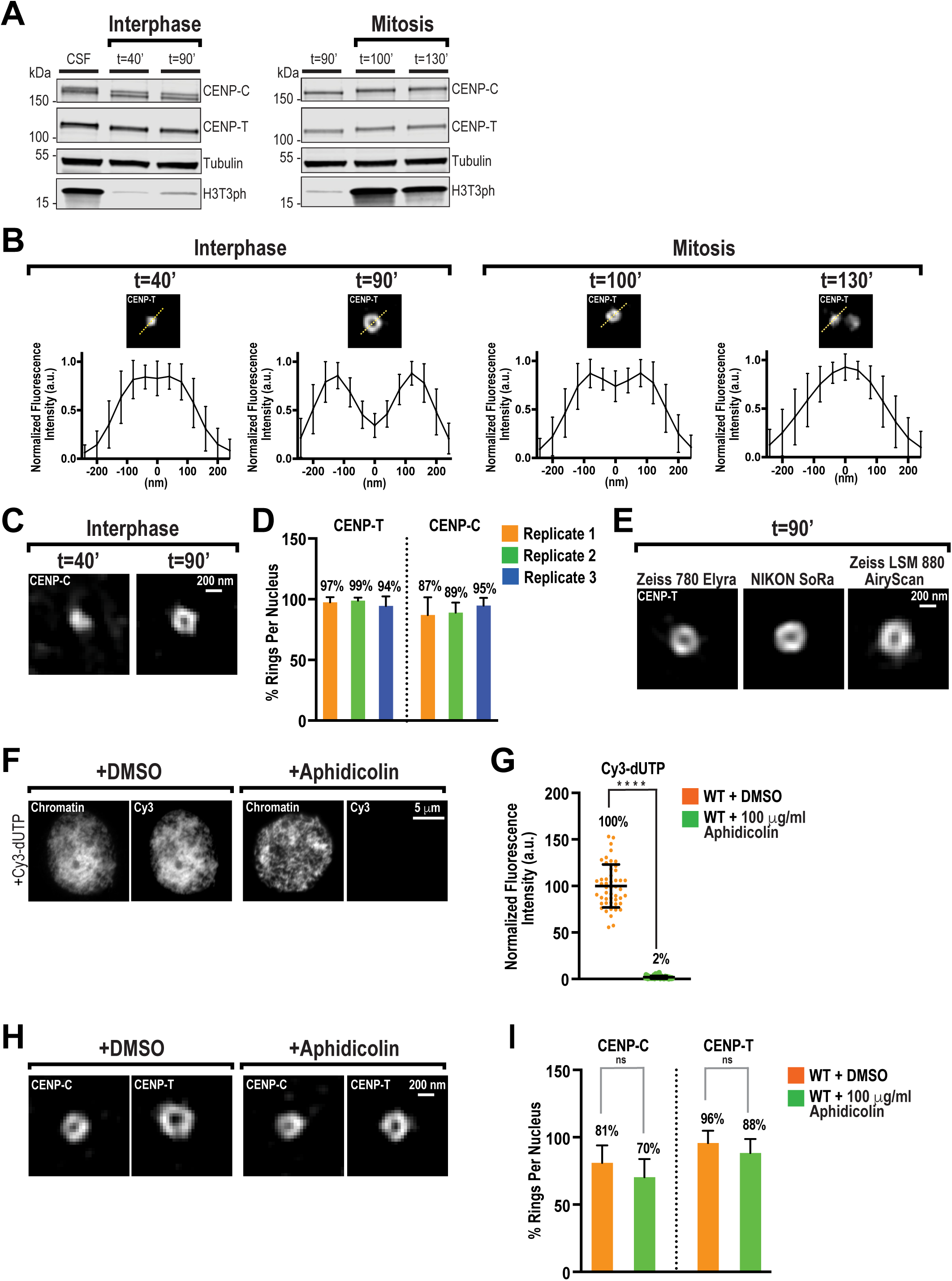
Centromeric chromatin architecture changes during the cell cycle but is independent of DNA replication. (**A**) Western blot for CENP-C, CENP-T, Tubulin, and Histone H3 phosphorylation (H3T3ph) for samples shown in Fig. 1A. CSF (cytostatic factor) *Xenopus* egg extracts are arrested in M phase without prior cycling through interphase. (**B**) Averaged and aligned linescan quantification of CENP-T fluorescence at centromeres from Fig. 1A. Each individual trace (not shown) was aligned by its central minimum or maximum and normalized to the maximum intensity, and the average trace plotted on this scale. Insets illustrate the line scanning method. n=75 kinetochores for each condition. Error bars represent SD. (**C**) Representative 3D-SIM images of CENP-C on interphase nuclei at *t*=40 and 90’ assembled from WT extract. Scale bar is 200 nm. (**D**) Percentage of observable ring-like structures per nucleus for CENP-C and CENP-T from three independent experiments. n=10 nuclei per condition, and n≥145 centromeres total. Error bars represent SD. (**E**) Representative super-resolution images of CENP-T on interphase nuclei at *t*=90, in WT extract acquired on the indicated microscope platforms. Scale bar is 200 nm. (**F**) Representative fluorescence images of chromatin (DAPI) and Cy3-dUTP on interphase nuclei, at *t*=90’, assembled from extract treated with DMSO or Aphidicolin. Scale bar is 5 µm. (**G**) Quantification of fluorescence intensity of Cy3-dUTP fluorescence shown in (F), normalized to DMSO. n = 50 nuclei per condition. A.U., arbitrary units. Error bars represent SD, ****, *P* < 0.0001. (**H**) Representative 3D-SIM images of CENP-C and CENP-T on interphase nuclei, at *t*=90’, assembled from extract treated with DMSO or Aphidicolin. Scale bar is 200 nm. (**I**) Quantification of percentage of CENP-C and CENP-T rings per nucleus following treatment with DMSO or Aphidicolin, shown in (H). n=10 nuclei per condition, and n≥145 centromeres total. Error bars represent SD. NS = Not significant

**Fig. S2.**
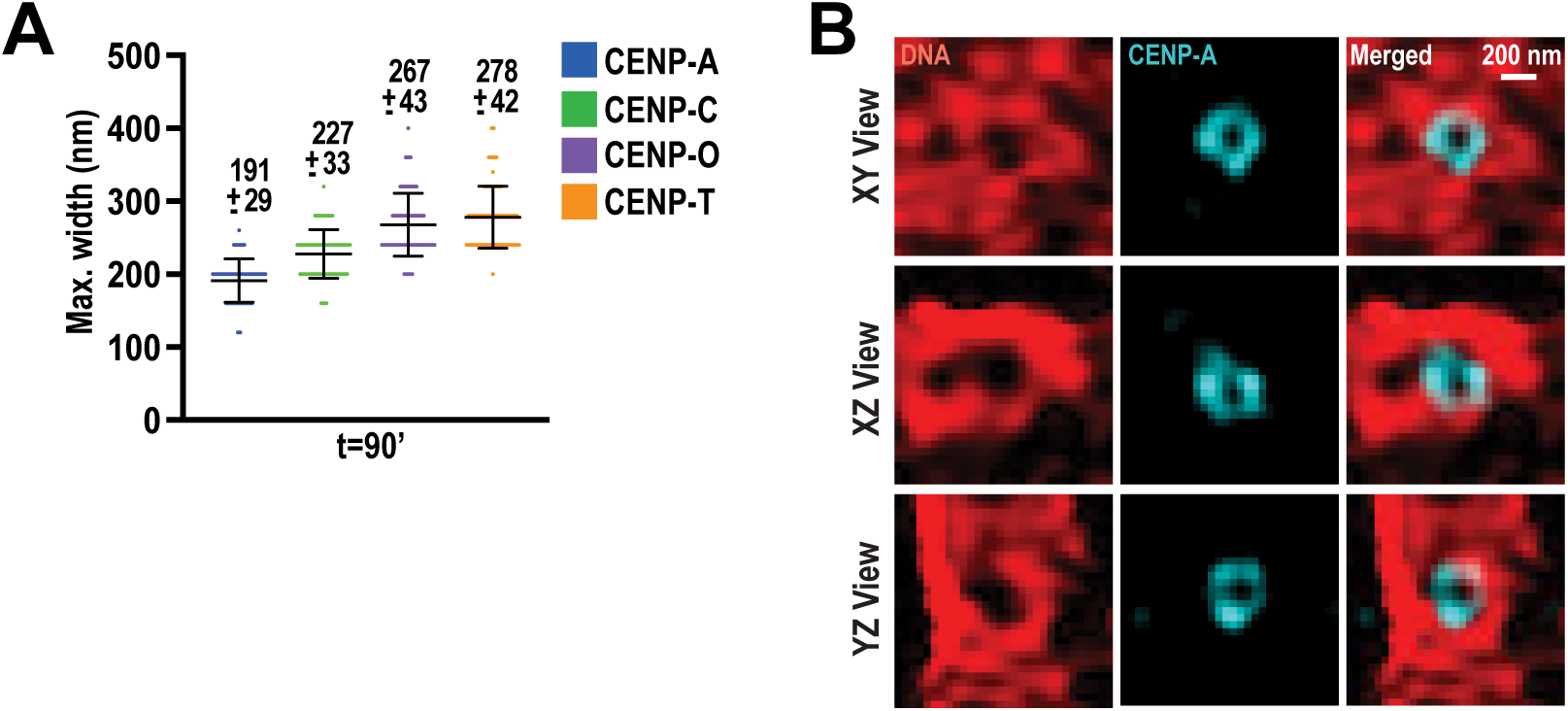
3D-SIM and deep learning analysis of CCAN and CENP-A structures. **(A)** Average peak-to-peak distances of CENP-A, CENP-C, CENP-O, and CENP-T signals from linescans of 3D-SIM images of interphase nuclei at t=90’, with representative images shown in Fig. 1B. n = 75 centromeres or kinetochores were quantified. Error bars represent SD. (**B**) Axial resolution enhancement of 3D-SIM images of CENP-A (cyan) and DNA (DAPI; red) from deep learning prediction. Representative XY, XZ, and YZ views of CENP-A and DNA are shown. Scale bar is 200 nm.

**Fig. S3.**
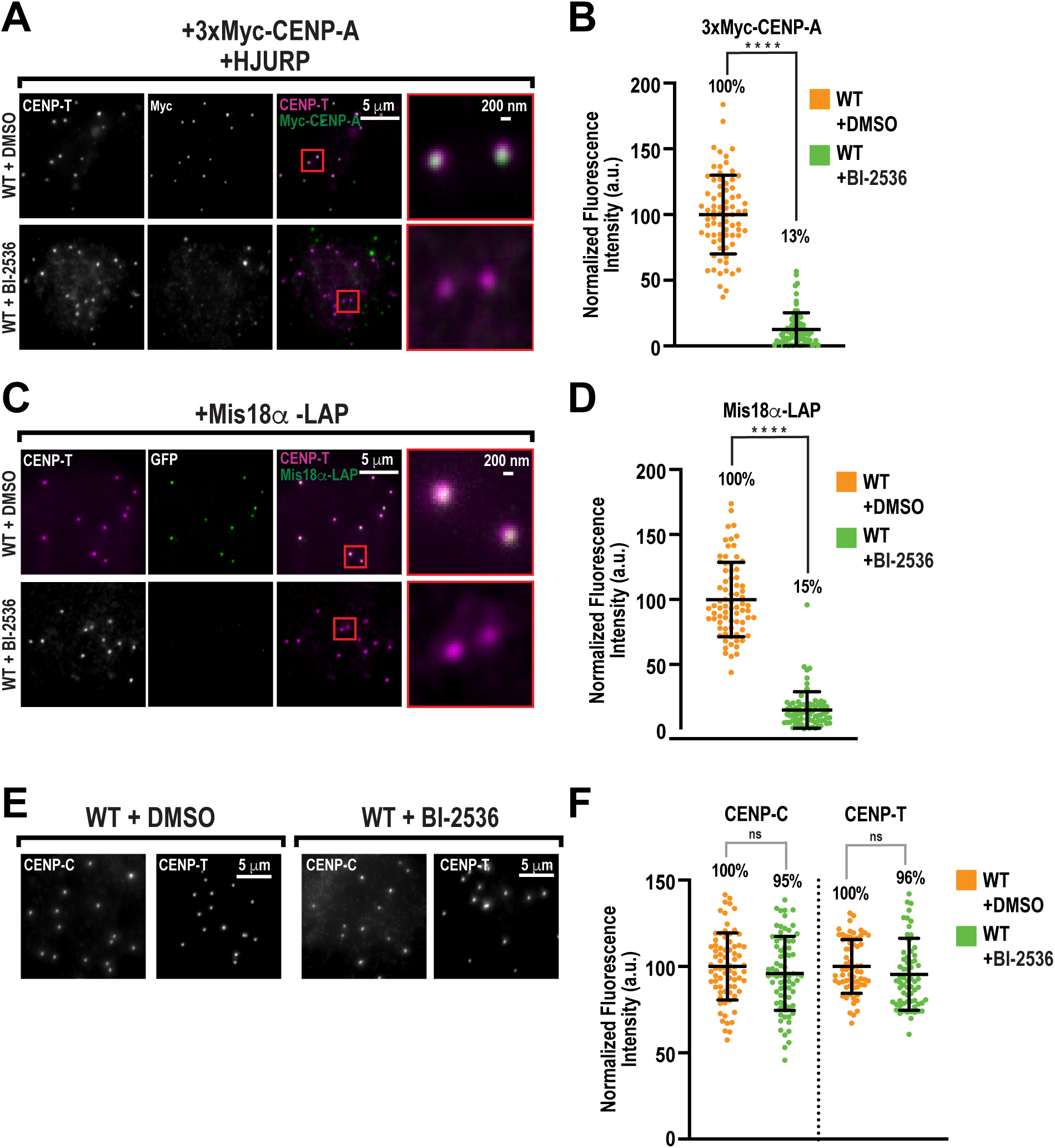
Inhibition of Plk1 activity blocks new CENP-A assembly in *Xenopus* egg extracts. **(A)** Representative immunofluorescence images of CENP-T (magenta) and 3XMyc-CENP-A (green) on interphase nuclei, at t=90’, assembled from extracts containing 3X-Myc-CENP-A and HJURP mRNA and treated with DMSO or BI-2536. Higher magnification views of the red boxed regions are shown in the last column. Scale bar is 5 µm, except for higher magnification views (200 nm). (**B**) Quantification of fluorescence intensity of 3XMyc-CENP-A shown in (A) normalized to DMSO. n = 75 kinetochores per condition. A.U., arbitrary units. Error bars represent SD. (**C**) Representative immunofluorescence images of CENP-T (magenta) and Mis18α-GFP (green) on interphase nuclei, at t=90’, assembled from extracts containing *in vitro* translated Mis18α-LAP treated with either DMSO or BI-2536. Higher magnification views of the red boxed regions are shown in the last column. Scale bar is 5 µm, except for higher magnification views (200 nm). (**D**) Quantification of fluorescence intensity of Mis18α-GFP shown in (C) normalized to DMSO. n = 75 kinetochores per condition. A.U., arbitrary units. Error bars represent SD. (**E**) Representative immunofluorescence images of CENP-C and CENP-T on interphase nuclei, at *t*=90’, assembled from extract treated with DMSO or BI-2536. Scale bar is 5 µm. (**F**) Quantification of fluorescence intensity of CENP-C and CENP-T shown in (E), normalized to DMSO. n=75 kinetochores per condition. A.U., arbitrary units. Error bars represent SD.

**Fig. S4.**
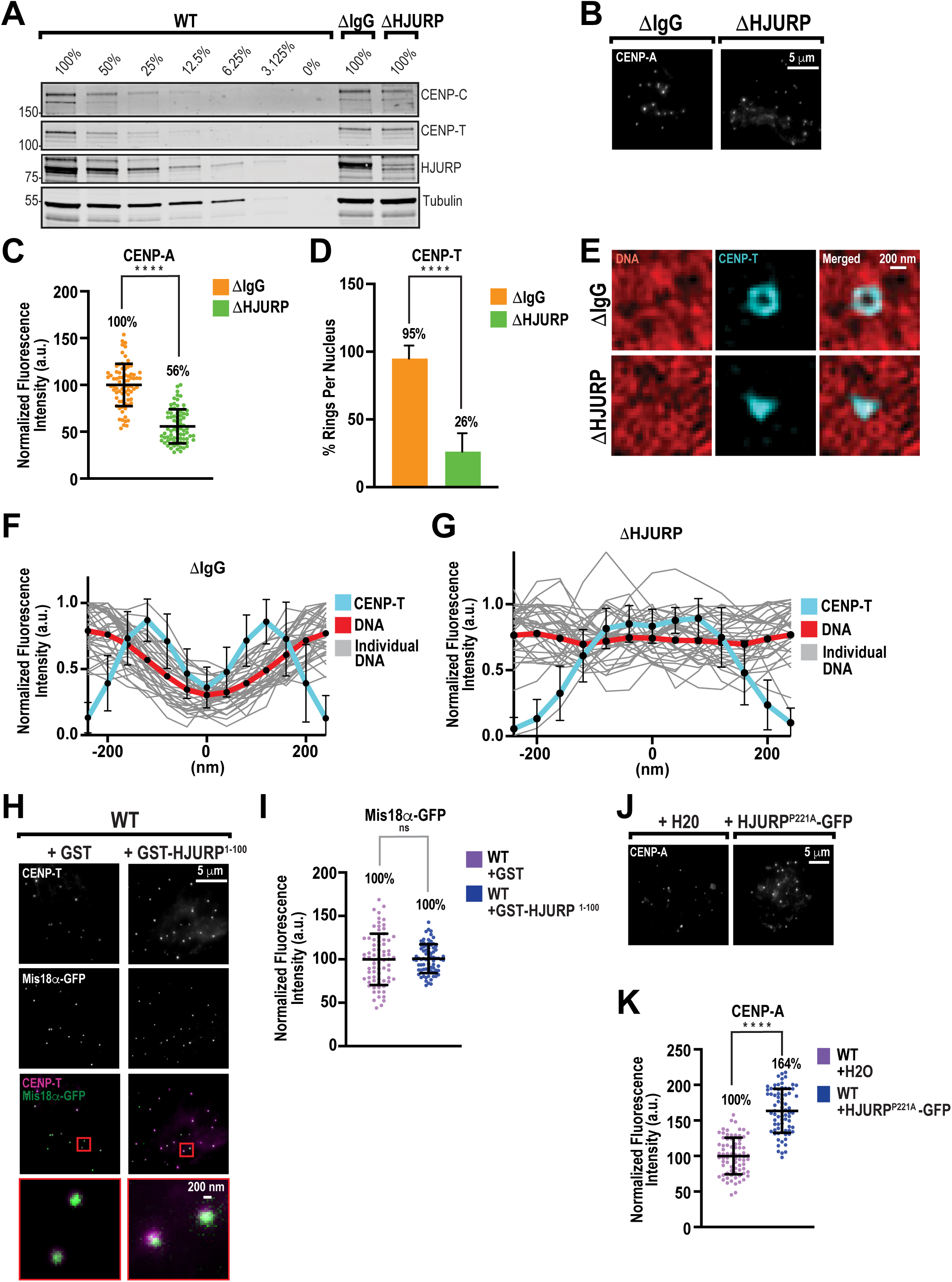
HJURP, but not its role in CENP-A assembly, is required for centromere organization in interphase. (**A**) Western blot for CENP-C, CENP-T, HJURP, and Tubulin in WT, ΔIgG, and ΔHJURP extracts from the samples shown in Fig. 3A. (**B**) Representative immunofluorescence images of CENP-A on interphase nuclei, at *t*=90’, assembled from ΔIgG and ΔHJURP extracts. Scale bar is 5 µm. (**C**) Quantification of fluorescence intensity of CENP-A shown in (B), normalized to ΔIgG. n=75 centromeres per condition. A.U., arbitrary units. Error bars represent SD, ****, *P* < 0.0001. (**D**) Percentage of observable CENP-T ring-like structures per nucleus for conditions in Fig. 3A. n=10 nuclei per condition, and n≥145 centromeres total. Error bars represent SD, ****, *P* < 0.0001. (**E**) Representative 3D-SIM images of CENP-T (cyan) and DNA (DAPI; red) on interphase nuclei, at *t*=90’, assembled in ΔIgG or ΔHJURP extracts. Scale bar is 200 nm. **(F**) Linescan quantification of CENP-T and DNA fluorescence at kinetochores in ΔIgG condition from (E). Bold lines indicate average intensity, and thin gray lines indicate individual traces of DNA fluorescence. Each trace was normalized to the maximum intensity and aligned to the central minimum of CENP-T average intensity. n=35 kinetochores. Error bars represent standard deviation (SD). (**G**) Linescan quantification of CENP-T and DNA fluorescence at kinetochores in ΔHJURP condition from (E). Bold lines indicate average intensity, and thin gray lines indicate individual traces of DNA fluorescence. Each trace was normalized to the maximum intensity and aligned to the central maximum of CENP-T average intensity. n=35 kinetochores. Error bars represent standard deviation (SD). (**H**) Representative immunofluorescence images of CENP-T (magenta) and Mis18α-GFP (green) on interphase nuclei, at t=90’, assembled from extracts containing *in vitro* translated Mis18α-GFP treated with GST or GST-HJURP^1-100^. Higher magnification views of the red boxed regions are shown in the last column. Scale bar is 5 µm, except for higher magnification views (200 nm). (**I**) Quantification of fluorescence intensity of Mis18α-GFP shown in (H) normalized to GST. n= 75 kinetochores per condition. A.U., arbitrary units. Error bars represent SD. (**J**) Representative immunofluorescence images of CENP-A on interphase nuclei, at *t*=90’, assembled from extracts containing water (H2O) or HJURP^P221A^-GFP mRNA. Scale bar is 5 µm. (**K**) Quantification of fluorescence intensity of CENP-A shown in (J), normalized to H2O. n=75 centromeres per condition. A.U., arbitrary units. Error bars represent SD, ****, *P* < 0.0001.

**Fig. S5.**
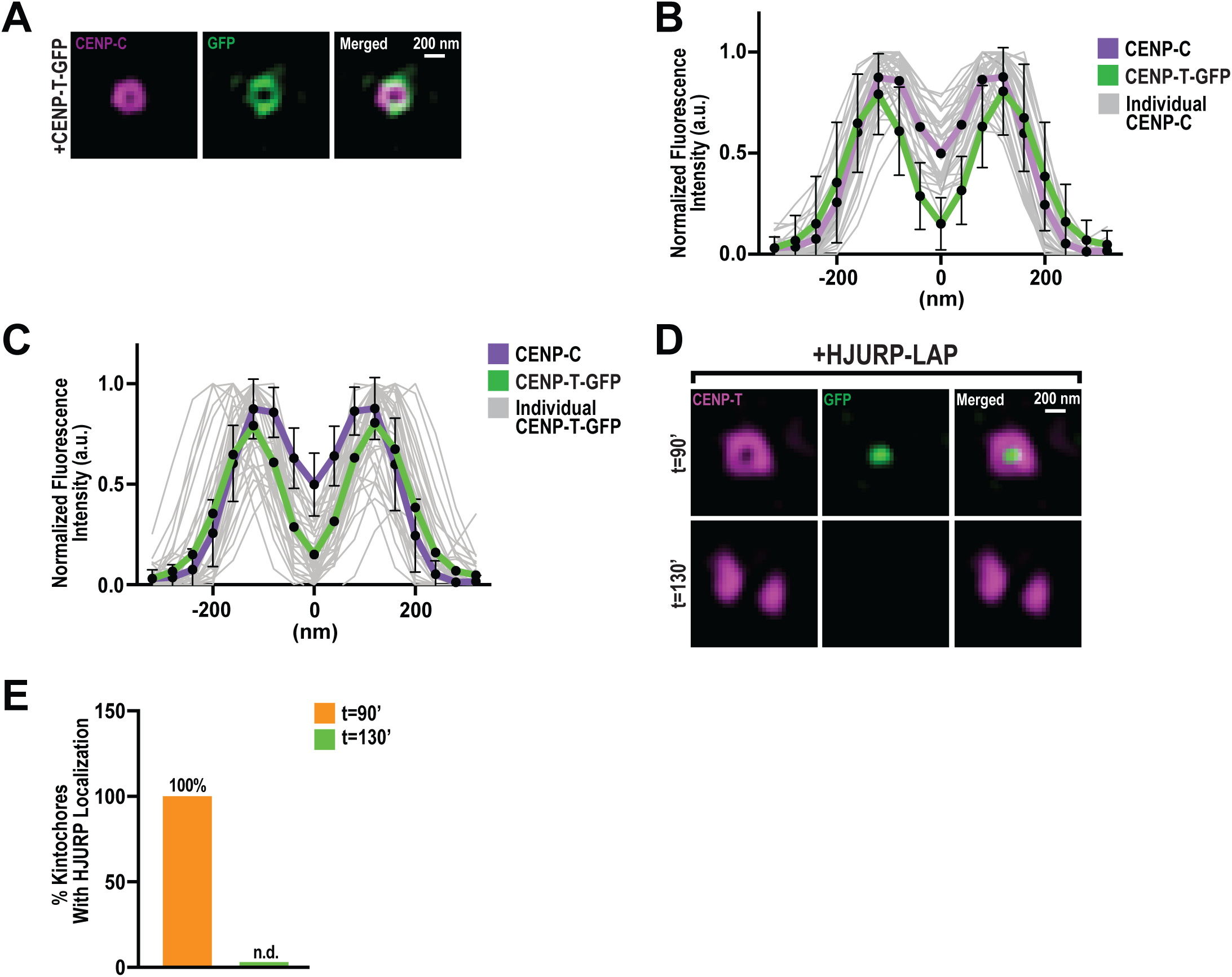
HJURP is evicted in mitosis, which correlates with the collapse of centromere organization. (**A**) Representative 3D-SIM images of CENP-C (magenta) co-stained with GFP (green) on interphase nuclei, at *t*=90’, assembled from extracts containing *in vitro* translated CENP-T-GFP. Scale bar is 200 nm. (**B**) Line scan quantification of CENP-C and CENP-T-GFP fluorescence at centromeres from (A). Bold lines indicate average intensity, and thin gray lines indicate individual traces of CENP-C fluorescence. Each trace was aligned by the central minimum and normalized to the maximum intensity. n=75 kinetochores. Error bars represent SD. (**C**) Line scan quantification of CENP-C and CENP-T-GFP fluorescence at centromeres from (A). Bold lines indicate average intensity, and thin gray lines indicate individual traces of CENP-T-GFP fluorescence. Each trace was aligned by the central minimum and normalized to the maximum intensity. n=75 kinetochores. Error bars represent SD. (**D**) Representative 3D-SIM images of CENP-T (magenta) and HJURP-GFP (green) on interphase nuclei and mitotic chromosomes at *t* = 90’ (interphase) and 130’(metaphase), assembled from WT extract containing *in vitro* translated HJURP-GFP. (**E**) Percentage of centromeres with HJURP present on interphase nuclei and mitotic chromosomes shown in (D). n=75 kinetochores in each condition.

**Fig. S6.**
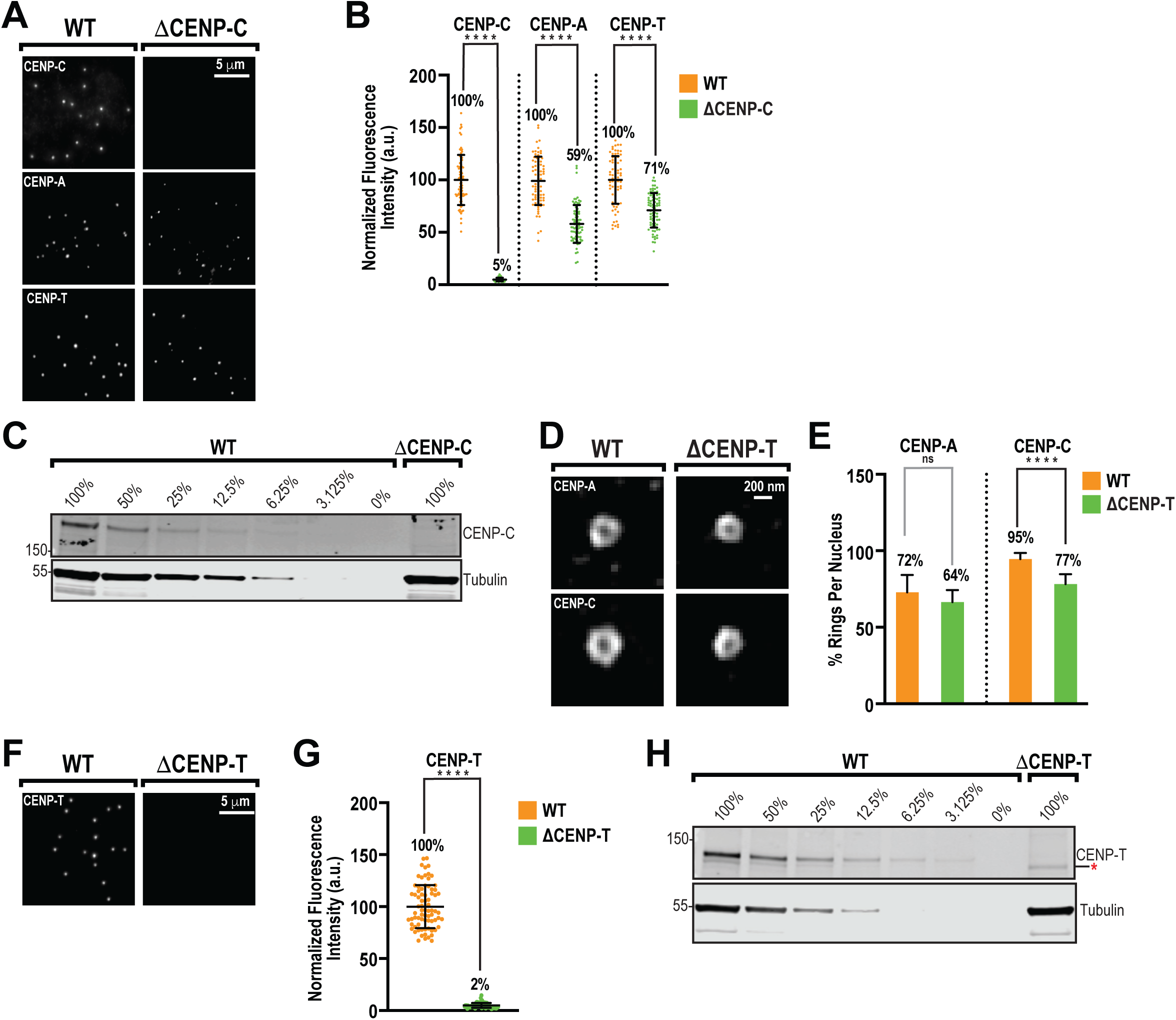
CENP-C, but not CENP-T, is required for centromere architecture in interphase. **(A)** Representative immunofluorescence images of CENP-C, CENP-A, and CENP-T on interphase nuclei, at *t*=90’, assembled from WT and CENP-C depleted (ΔCENP-C) extracts for the samples shown in Fig. 4D. Scale bar is 5 µm. (**B**) Quantification of fluorescence intensity of CENP-C, CENP-A, and CENP-T shown in (A) normalized to WT. n = 75 centromeres or kinetochores per condition. A.U., arbitrary units. Error bars represent SD, ****, *P* < 0.0001. (**C**) Western blot for CENP-C and Tubulin in WT and ΔCENP-C extracts from the samples shown in (A) and Fig. 4D. (**D**) Representative 3D-SIM images of CENP-A and CENP-C on interphase nuclei, at *t*=90’, assembled from WT and CENP-T depleted (ΔCENP-T) extracts. Scale bar is 200 nm. (**E**) Percentage of observable CENP-A or CENP-C ring-like structures per nucleus for conditions in (D). n=8 nuclei per condition, and n≥129 centromeres total. Error bars represent SD, ****, *P* < 0.0001. (**F**) Representative immunofluorescence images of CENP-T on interphase nuclei, at *t*=90’, assembled from WT and ΔCENP-T extracts. Scale bar is 5 µm. (**G**) Quantification of fluorescence intensity of CENP-T immunofluorescence from (F), normalized to WT. n = 75 kinetochores per condition. A.U., arbitrary units. Error bars represent SD, ****, *P* < 0.0001. (**H**) Western blot for CENP-T and Tubulin in WT and ΔCENP-T extracts from the samples shown in (D). Asterisk indicates a non-specific band.

